# POMC-specific knockdown of Tril reduces body adiposity and increases hypothalamic leptin responsiveness

**DOI:** 10.1101/2020.06.25.172379

**Authors:** Alexandre Moura-Assis, Pedro A. Nogueira, Jose C. de-Lima-Junior, Fernando M. Simabuco, Joana M. Gaspar, Jose Donato, Licio A. Velloso

**Affiliations:** Laboratory of Cell Signalling-Obesity and Comorbidities Research Center, University of Campinas, Campinas, Brazil; Multidisciplinary Laboratory of Food and Health (LABMAS), School of Applied Sciences (FCA), University of Campinas (UNICAMP), Limeira, São Paulo, Brazil; Department of Physiology and Biophysics, Institute of Biomedical Sciences, University of Sao Paulo, Sao Paulo, Brazil; The Laboratory of Cell Signalling belongs to the National Institute of Science and Technology on Neuroimmunomodulation (INCT-NIM)

## Abstract

In a public dataset of transcripts differentially expressed in selected neuronal subpopulations of the arcuate nucleus, we identified TLR4-interactor with leucine-rich repeats (Tril) as a potential candidate for mediating the harmful effects of a high-fat diet in proopiomelanocortin (POMC) neurons. The non-cell-specific inhibition of Tril in the arcuate nucleus resulted in reduced hypothalamic inflammation, protection against diet-induced obesity associated with increased whole-body energy expenditure and increased systemic glucose tolerance. The inhibition of Tril, specifically in POMC neurons, resulted in a trend for protection against diet-induced obesity, increased energy expenditure and increased hypothalamic sensitivity to leptin. Thus, Tril emerges as a new component of the complex mechanisms that promote hypothalamic dysfunction in experimental diet-induced obesity.

## Introduction

In diet-induced obesity (DIO), nutrients, particularly saturated fats, trigger a toll-like receptor-4 (TLR4)-dependent inflammatory response that affects neurons of the mediobasal hypothalamus (MBH) (1–3). Proopiomelanocortin (POMC) neurons are particularly sensitive to this inflammatory response undergoing apoptosis at higher rates than agouti-related peptide (AgRP) neurons (4–6). This can lead to an imbalance in the rate of anorexigenic/orexigenic neurons, potentially explaining the chronic and recurrent nature of obesity (6). Thus, defining the mechanisms that selectively act upon POMC neurons rendering them more sensitive to inflammatory insults could provide advances in the understanding of the hypothalamic pathophysiology of obesity.

Methodological progress in transcriptome analysis has refined the capacity of characterizing gene expression patterns in selected cellular populations (7). Using a cell-type specific transcriptome analysis of AgRP and POMC neurons of the MBH, Henry and coworkers (8) built a dataset of transcripts expressed in each of the main hypothalamic neuronal subpopulations involved in the control of food intake and energy expenditure. We employed this dataset in a search for transcripts encoding proteins related to the TLR4 signaling system that could be predominantly expressed in POMC neurons; we identified TLR4-interactor with leucine-rich repeats (Tril) as a candidate to mediate some of the harmful effects of a high-fat diet (HFD) in POMC neurons.

Tril is a single transmembrane spanning 89 kDa protein that contains 13 leucine-rich repeats and is highly expressed in the brain (9). It plays an important role mediating TLR4 signal transduction, and whole-body Tril knockout results in defective inflammatory cytokine production in response to LPS and *E. coli*, particularly in the central nervous system (CNS) (10). Here, we hypothesized that Tril could be involved in diet induced-hypothalamic inflammation and particularly in the regulation of POMC neurons. We show that knockdown of hypothalamic Tril protects mice from diet-induced body mass gain and systemic glucose intolerance.

## Results

### Hypothalamic Tril expression is increased in diet-induced obesity

In POMC (Fig. 1A) and AgRP (Fig. 2B) reporter mice, Tril was predominantly detected in POMC neurons; Tril was also detected in some Iba1 expressing cells (Fig. 1C). In outbred obesity-prone Swiss mice (Fig. 1D), the hypothalamic (Fig. 1E) but not hippocampal (Fig. 1F) expression of Tril underwent increase 1 and 2 weeks after the introduction of a HFD. This was accompanied, in the first week, by the increased expression of hypothalamic Tlr4 (Fig. 1G). In isogenic C57BL/6J mice, the hypothalamic expression of Tril was increased after refeeding in mice fed a HFD (Fig. 1H) but not in mice fed chow (Fig. 1I).

**Figure 1.**
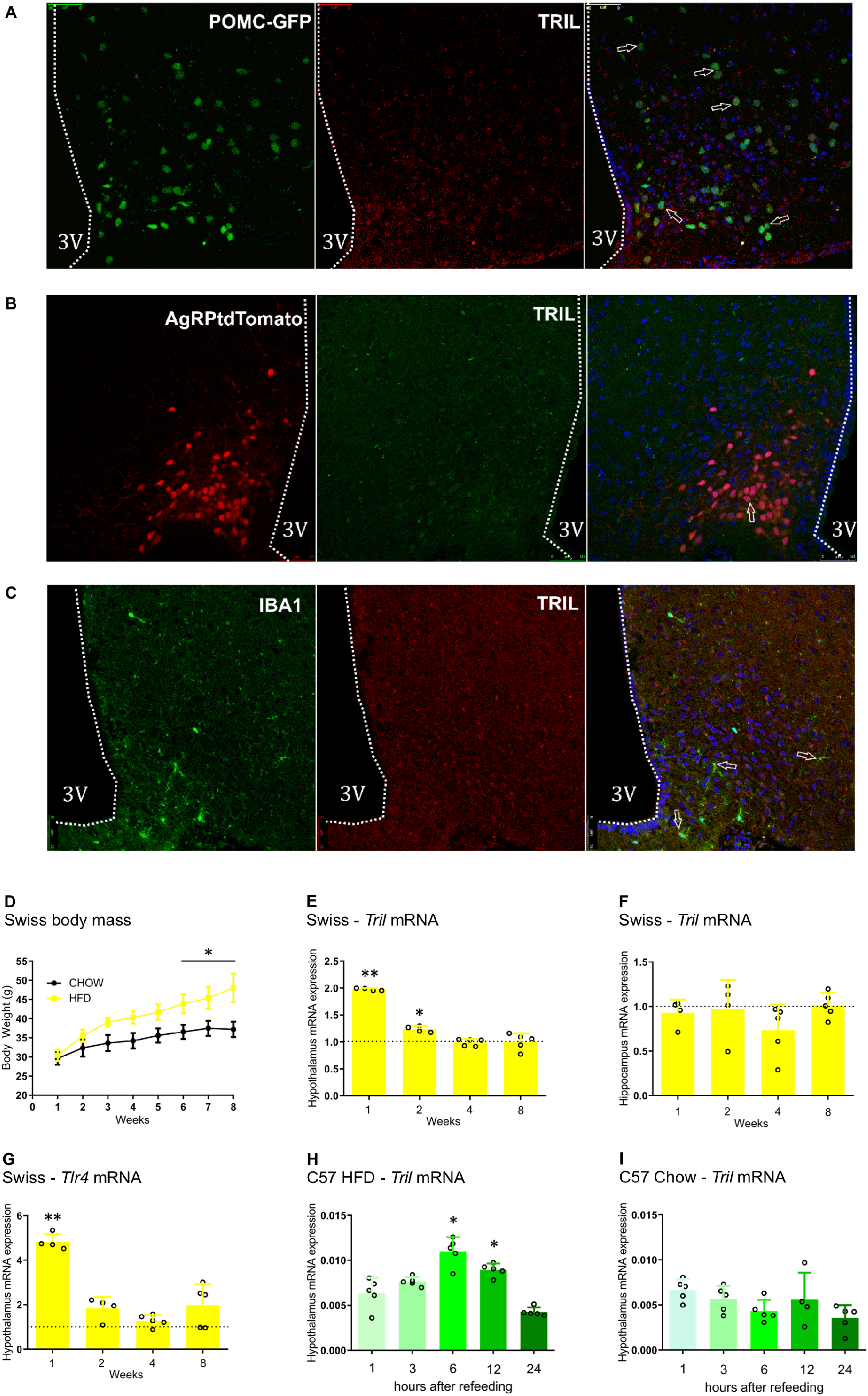
The impact of a high-fat diet on the expression of Tril. The hypothalamic expression of Tril was determined using immunofluorescence staining in sections obtained from chow fed POMC-GFP (A), AgRPtdTomato (B) and C57BL/6J (C) mice; in A and B, POMC and AgRP neurons were detected as expressing the respective endogenous fluorescent peptides, whereas in C, Iba1-expressing cells were detected using a specific antibody. In A–C, Tril was detected using a specific antibody; the arrows indicate cells co-expressing target peptides. In D–G, Swiss mice were fed chow or a high-fat diet (HFD) for 1 to 8 weeks; body mass is shown in D; transcript expressions of Tril (E, hypothalamus; F, hippocampus) and Tlr4 (G, hypothalamus) in mice fed a HFD are plotted as normalized for the respective expressions in mice fed chow (broken line). C57BL/6J mice fed a HFD (H) or chow (I) were submitted to an overnight fast; diets were offered and hypothalami were extracted after 1, 6, 12 or 24 h for determination of Tril transcript expression. In A–C, images are representative of three independent experiments. In D–G, n=5; *p<0.05 and **p<0.01 vs. respective control (chow). In H and I, n=5; *p<0.05 vs. 1 h.

**Figure 2.**
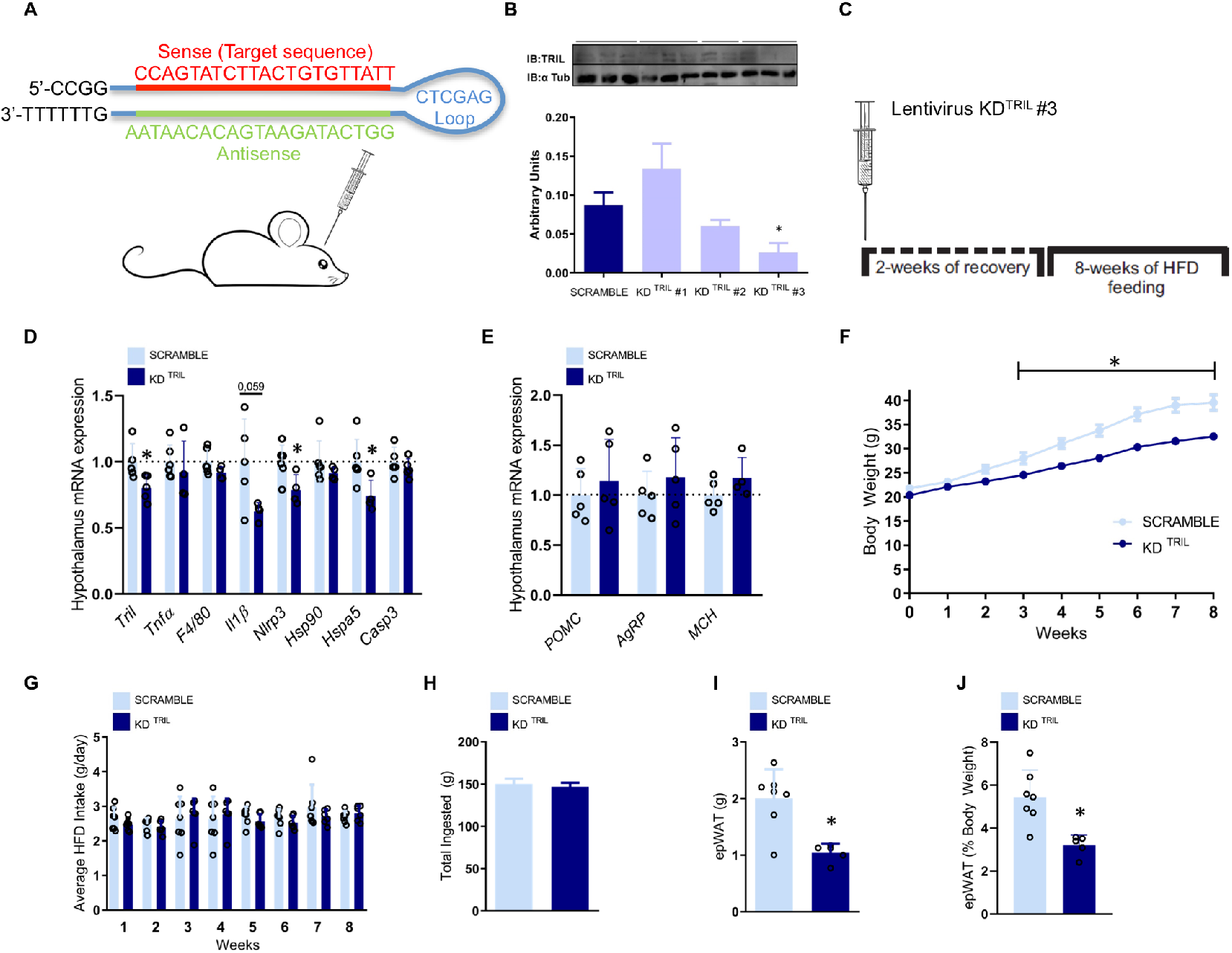
The inhibition of hypothalamic Tril affects body mass and adiposity. C57BL/6J mice were submitted to an intracerebroventricular injection of a lentivirus carrying a scramble or one out of three different sequences for targeting Tril (KD^TRIL^#1, KD^TRIL^#2 or KD^TRIL^#3) (A and B); immunoblot of hypothalamic extracts was employed to determine Tril expression in each experimental group (B); the sequence of the selected inhibitory sequence, KD^TRIL^#3, is depicted in A. The protocol employed in the experiments is depicted in C. The hypothalamic transcript expression of inflammatory and apoptotic genes (D) and neurotransmitter genes (E) was determined at the end of the experimental period. Body mass (F) and food intake (G and H) were determined during the experimental period. Total and relative (J) epididymal fat mass were determined at the end of the experimental period. In B, n=4; in D–J, n=5–7. In all experiments *p<0.05 vs. scramble. epWAT, epididymal white adipose tissue; HFD, high-fat diet; KD, knockdown.

### Inhibition of hypothalamic Tril protects from diet-induced obesity

Three distinct lentiviral clones encoding mouse shRNA to knockdown Tril were tested in the MBH (Fig. 2A and 2B); the best result was obtained employing lentivirus clone #3 (the sequence of lentivirus #3 is depicted in Fig. 2A and Tril protein inhibition in Figure 2B). The protocol (using lentivirus #3) was designed to evaluate whether the knockdown of hypothalamic Tril could prevent the effect of a HFD on metabolic parameters, since the introduction of a HFD occurred after the injection of the lentivirus particles (Fig. 2C). The inhibition of hypothalamic Tril resulted in the reduction of transcript expression of hypothalamic Il1b, Nlrp3 and Hspa5 (Fig. 2D); no changes were detected in baseline expression of transcripts encoding for Pomc, Agrp and Mch (Fig. 2E). The inhibition of hypothalamic Tril resulted in reduced body mass gain (Fig. 2F) and no change in caloric intake (Fig. 2G and 2H); in addition, inhibition of hypothalamic Tril resulted in reduced absolute (Fig. 2I) and relative (Fig. 2J) epididymal white adipose tissue mass.

### Inhibition of hypothalamic Tril increases whole body energy expenditure and improves systemic glucose tolerance

In mice under hypothalamic inhibition of Tril and fed a HFD (as depicted in the experimental protocol in Fig. 2C), there was an increase in whole body energy expenditure during the dark cycle (Fig. 3A–3D). This was accompanied by reduced absolute (Fig. 3E) and relative (Fig. 3F) mass of the interscapular brown adipose tissue (iBAT) but no changes in the temperature (Fig. 3G–3I) of iBAT and expression of iBAT transcripts encoding proteins involved in thermogenesis (Fig. 3J). Moreover, the inhibition of hypothalamic Tril improved systemic glucose tolerance (Fig. 3K and 3L), promoted a trend to increase systemic insulin sensitivity (Fig. 3M) and reduced liver steatosis (Fig. 3N), which was accompanied by reduced hepatic expression of transcripts encoding for Scd1 and CD36 (Fig. 3O).

**Figure 3.**
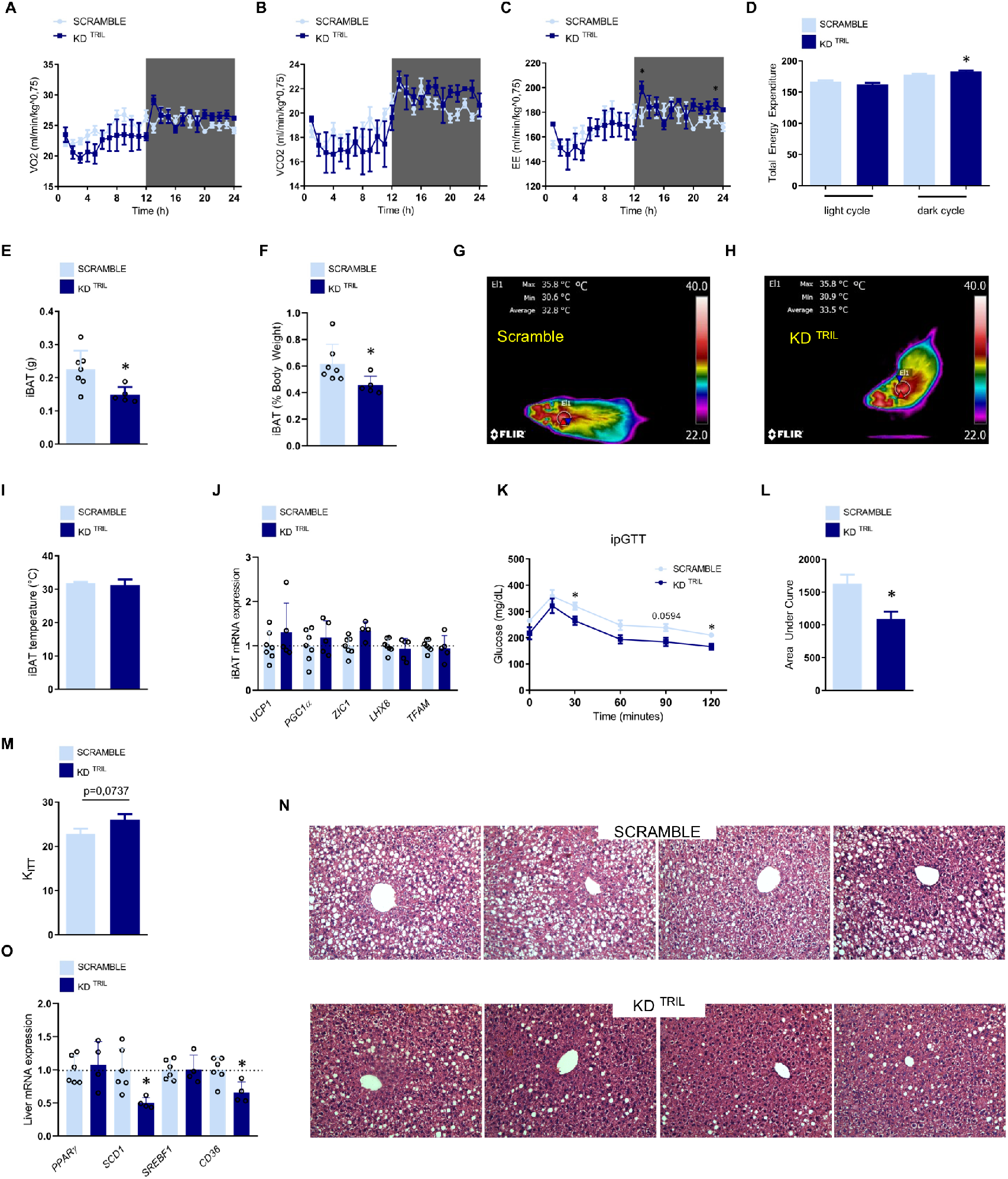
The inhibition of hypothalamic Tril affects energy expenditure and systemic metabolic parameters. C57BL/6J mice were submitted to an intracerebroventricular injection of a lentivirus carrying a scramble or an anti-Tril sequence (sequence KD^TRIL^#3) employing the same experimental protocol as depicted in Fig. 2C. At the end of the experimental period, mice were submitted to determination of O_2_ consumption (A), CO_2_ production (B), energy expenditure (C and D), determination of interscapular brown adipose tissue total (E) and relative (F) mass, determination of interscapular temperature (G–I), determination of interscapular brown adipose transcript expression of thermogenic genes, determination of whole-body glucose tolerance by means of an intraperitoneal glucose tolerance test (K, time course of blood glucose levels; L, area under the curve of blood glucose levels) and determination of whole body insulin sensitivity by means of an insulin tolerance test (M, constant of blood glucose decay during the insulin tolerance test). The liver was extracted for histological examination (N) and also for determination of transcript expression of genes involved in lipogenesis or lipid uptake (O). In A–M and O, n=5–7; *p<0.05 vs. scramble. In N, images are representative of four independent experiments. EE, energy expenditure; iBAT, interscapular brown adipose tissue; ipGTT, intraperitoneal glucose tolerance test; Kitt, constant of blood glucose decay during the insulin tolerance test.

### POMC-specific knockdown of Tril reduces body adiposity and increases hypothalamic responsiveness to leptin

A Cre-dependent rAAV was designed to selectively knockdown Tril in POMC neurons of the MBH (Fig. 4A and 4B). Since the introduction of a HFD occurred after the injection of rAAV, this protocol was designed to evaluate whether the knockdown of Tril in POMC neurons could prevent the effect of a HFD on metabolic parameters (Fig. 4C). This approach resulted in a trend to overcome HFD-induced body mass gain (Fig. 4D and 4E) and a significant reduction of fat (Fig. 4F) but not lean mass (Fig. 4G). In addition, there were trends to reduce absolute (Fig. 4H) and relative (Fig. 4I) epididymal fat mass and significant reductions of epididymal adipose tissue expression of the inflammatory transcripts Il1b and Tnfα (Fig. 4J). The knockdown of Tril in POMC neurons resulted in a trend to reduce cumulative food intake throughout the experimental period (Fig. 5A) and a significant reduction of food intake acutely after a period of prolonged fasting (Fig. 5B). This was accompanied by increased leptin-induced activation of STAT3 in the arcuate nucleus (Fig. 5C and 5E) but not in the retrochiasmatic hypothalamus (Fig. 5D). Inhibiting Tril in POMC neurons promoted no modification in the density of αMSH (Fig. 5F, upper panels and Fig. 5G) and AgRP (Fig. 5F, lower panels and Fig. 5H) projections to the paraventricular hypothalamus.

**Figure 4.**
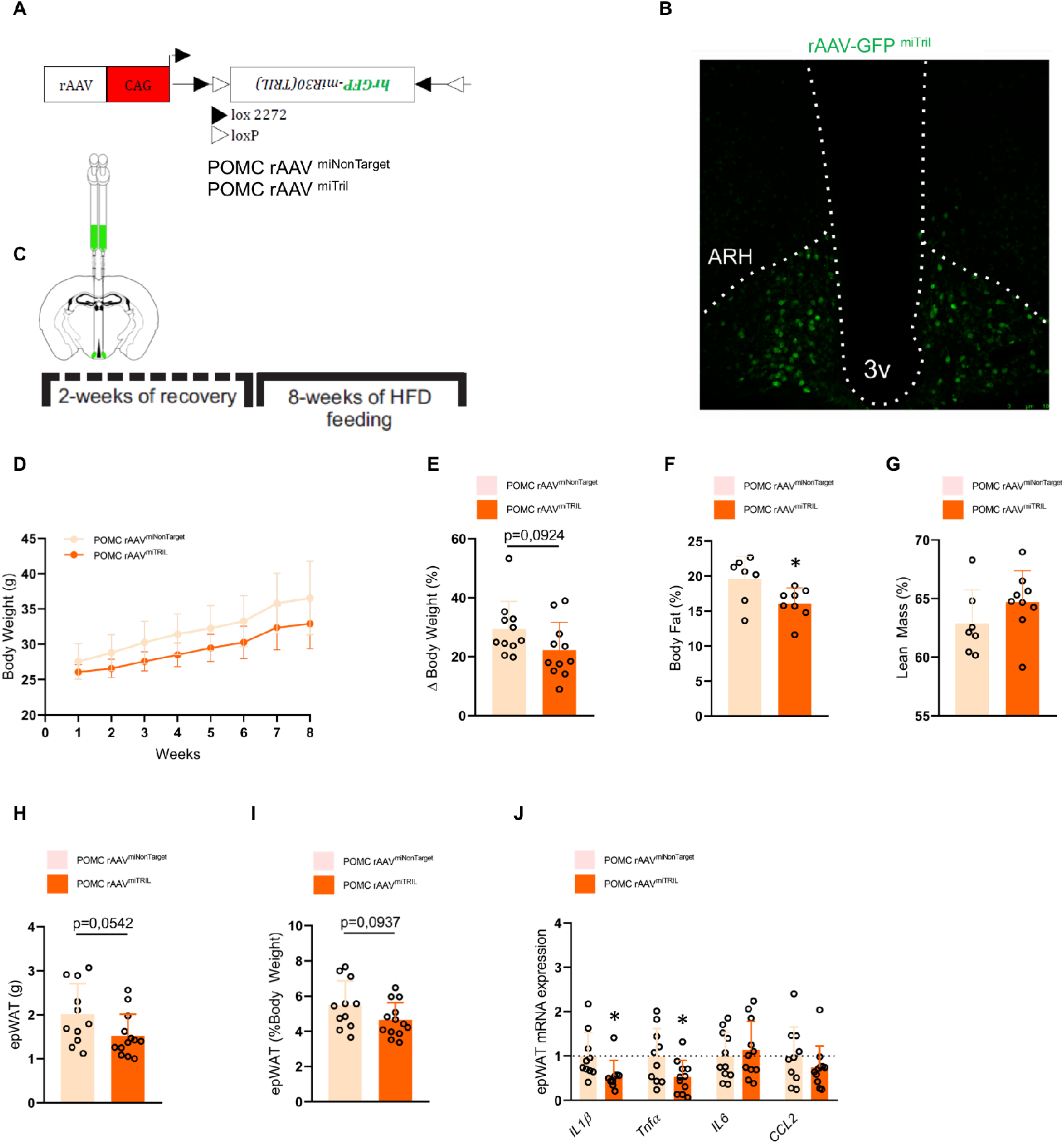
POMC-specific knockdown of Tril reduces body fat. POMC-Cre mice were submitted to an intracerebroventricular injection with Cre-dependent AAV-FLEX-EGFP-mir30 carrying either a non-target sequence (rAAVshNonTarget) or a Tril targeting sequence (rAAVshTril) (A). Evidence for the efficiency of the adenoviral incorporation into cells of the arcuate nucleus (ARH) was obtained by immunofluorescence evaluation of sections of the mediobasal hypothalamus (B). The protocol employed in the experiments is depicted in C. Body mass (D and E) was determined throughout the experimental period. Relative fat (F) and lean (G) mass as well as the absolute (H) and relative (I) epididymal fat mass were determined at the end of the experimental period. The expressions of inflammatory genes were determined in the epididymal adipose tissue at the end of the experimental period (J). In D–J, n=6–14; *p<0.05 vs. shNonTarget.

**Figure 5.**
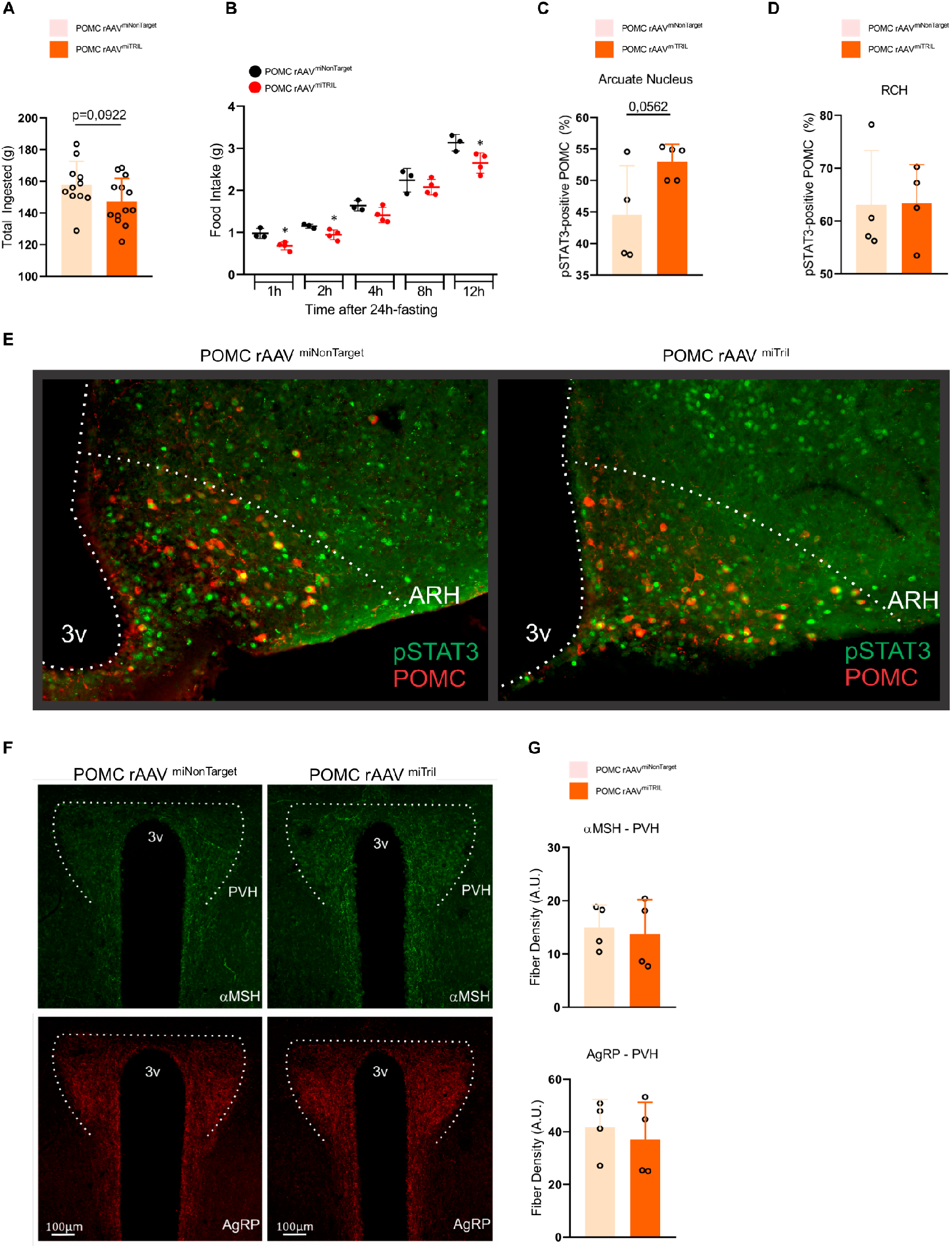
Inhibition of Tril in POMC neurons improves leptin sensitivity. POMC-Cre mice were submitted to an intracerebroventricular injection with Cre-dependent AAV-FLEX-EGFP-mir30 carrying either a non-target sequence (rAAVshNonTarget) or a Tril targeting sequence (rAAVshTril) and then submitted to the same protocol as in Fig. 4C. The cumulative consumption of diet was determined throughout the experimental period (A). At the end of the experimental period, mice were submitted to a determination of spontaneous food intake after a period of 24 h fasting (B). Leptin-induced phosphorylation of STAT3 was determined by calculating the proportion of phospho-STAT3 per POMC-positive cells in the arcuate nucleus (C) and retrochiasmatic hypothalamus (D) employing immunofluorescence staining; an illustrative image obtained from the mediobasal hypothalamus is depicted in E. Immunofluorescence staining was employed to determine the density of aMSH (F, upper panels and G) and AgRP (F, lower panels and H) fiber projections to the paraventricular hypothalamus. In A, n=11–13; in B–H, n=4–5; *p<0.05 vs. shNonTarget.

### POMC-specific knockdown of Tril increases whole body energy expenditure and thermogenic gene expression in brown adipose tissue

In mice fed chow, the POMC-specific knockdown of Tril resulted in increased whole body energy expenditure, both in the light and dark cycles (Fig. 6A–6D). In mice fed a HFD (according to the protocol depicted in Fig. 4C), the inhibition of Tril, specifically in POMC neurons, resulted in a trend to increase energy expenditure during the dark cycle (Fig. 6E–6H) and a trend to reduce absolute brown adipose tissue (Fig. 6I) but not relative mass (Fig. 6J). Brown adipose tissue temperature was not modified (Fig. 6K–6M), but the expression of the thermogenic genes Lhx8 and Zic1 were increased in mice under the inhibition of Tril in POMC neurons (Fig. 6N). In addition, the inhibition of Tril in POMC neurons led to lower fasting blood glucose (Fig. 6O) but no change in whole body glucose tolerance (Fig. 6P).

**Figure 6.**
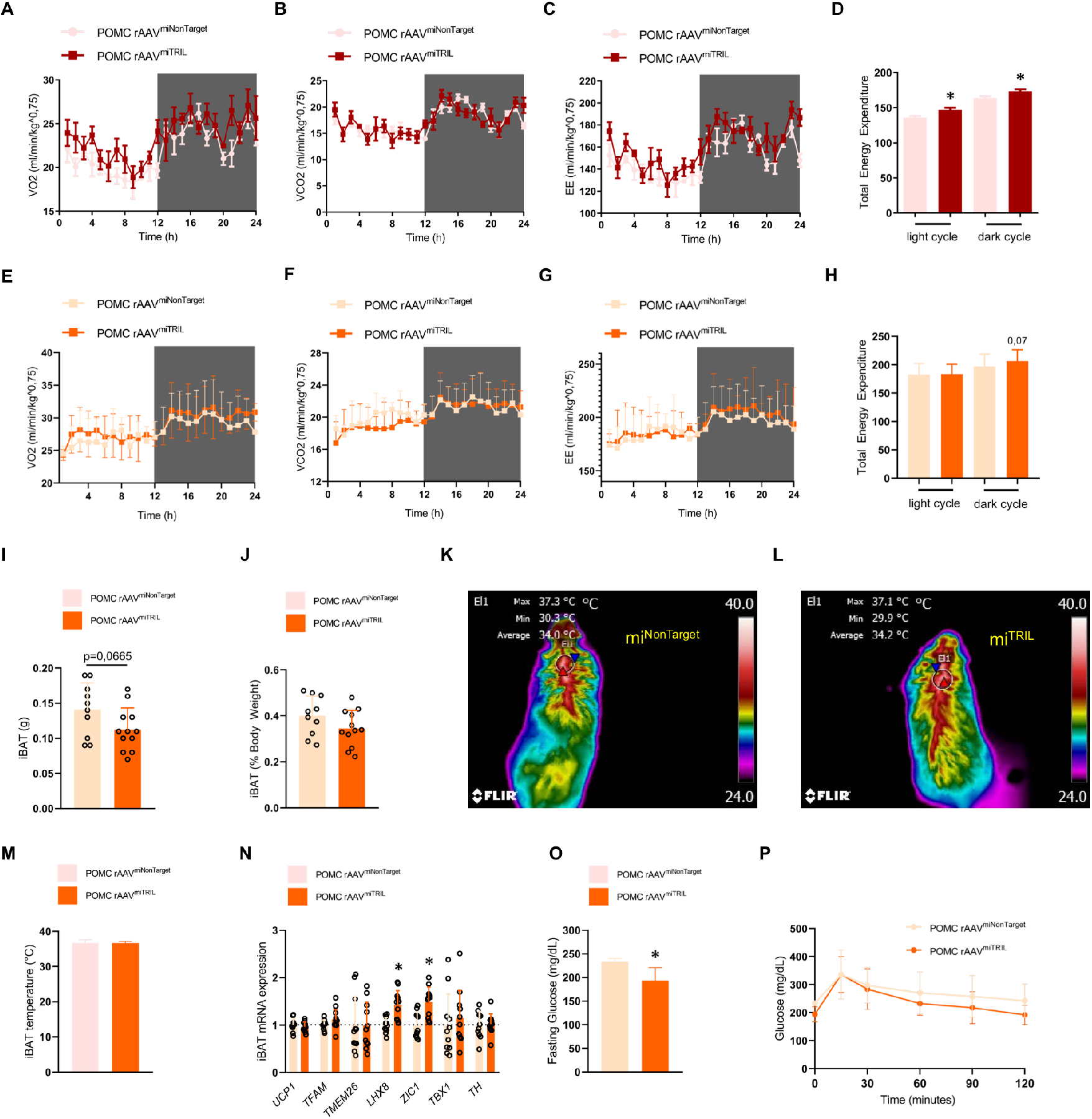
Inhibition of Tril in POMC neurons increases energy expenditure. In A–D, POMC-Cre mice were submitted to an intracerebroventricular injection with Cre-dependent AAV-FLEX-EGFP-mir30 carrying either a non-target sequence (rAAVshNonTarget) or a Tril targeting sequence (rAAVshTril) and then left to recover for 2 weeks followed by another 2 weeks fed on chow; at the end of the experimental period, mice were submitted to determination of O_2_ consumption (A), CO_2_ production (B) and energy expenditure (C and D). In E–P, POMC-Cre mice were submitted to an intracerebroventricular injection with Cre-dependent AAV-FLEX-EGFP-mir30 carrying either a non-target sequence (rAAVshNonTarget) or a Tril targeting sequence (rAAVshTril) and then submitted to the same protocol as in Fig. 4C. At the end of the experimental period, mice were submitted to determination of O_2_ consumption (E), CO_2_ production (F), energy expenditure (G and H), determination of interscapular brown adipose tissue total (I) and relative (J) mass, determination of interscapular temperature (K–M) and determination of interscapular brown adipose transcript expression of thermogenic genes (N). In addition, blood glucose levels were determined in fasting mice (O) and whole-body glucose tolerance was determined by means of an intraperitoneal glucose tolerance test (P). In A–H, K–M, O and P, n=5–7; in I, J and N, n=12–14. In all, *p<0.05 vs. shNonTarget.

### Inhibition of Tril in POMC neurons cannot revert the diet-induced obesity phenotype but increases energy expenditure and expression of thermogenic genes in brown adipose tissue

In all preceding experiments, the inhibition of Tril either in the whole hypothalamus or specifically in POMC neurons occurred 2 weeks before the introduction of a HFD; thus, the approaches aimed at preventing the harmful effects of the diet. In order to test the hypothesis that inhibition of Tril in POMC neurons could revert the metabolically adverse effects of long-term consumption of a HFD, mice were fed a HFD for 14 weeks and then injected with rAAV shTril. After 2 weeks of recovery, mice were maintained for another 8 weeks on the HFD and metabolic parameters were evaluated (Fig. 7A). As depicted in Fig. 7B and 7C, the inhibition of Tril in POMC neurons could neither revert obesity nor change caloric intake in this group of mice. In addition, there were no changes in epididymal fat mass (Fig. 7D and 7E), brown adipose tissue mass (Fig. 7F and 7G) and brown adipose tissue temperature (Fig. 7H–7J). Nevertheless, in mice submitted to the inhibition of Tril in POMC neurons, there was a trend to increase brown adipose tissue expression of Ucp1 and a significant increase in the expression of Tmem26 (Fig. 7K).

**Figure 7.**
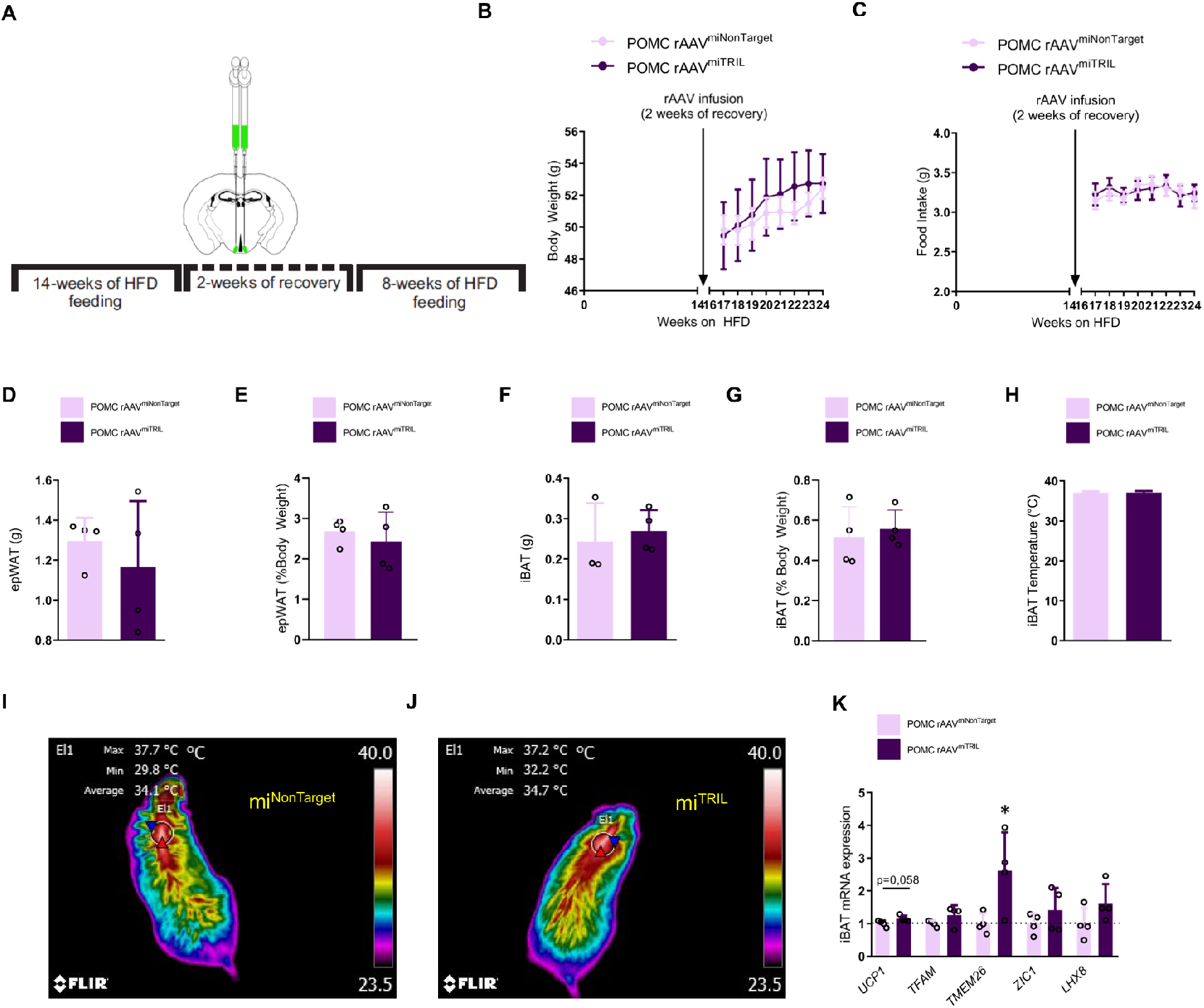
Inhibition of Tril in POMC neurons in mice submitted to long-term feeding on a high-fat diet fails to revert the obese phenotype. POMC-Cre mice were fed for 14 weeks on a high-fat diet and then submitted to an intracerebroventricular injection with Cre-dependent AAV-FLEX-EGFP-mir30 carrying either a non-target sequence (rAAVshNonTarget) or a Tril targeting sequence (rAAVshTril); after 2 weeks of recovery, mice were fed a high-fat diet for another 8 weeks (experimental protocol depicted in A). Body mass (B) and food intake (C) were recorded throughout the experimental period. Relative fat (F) and lean (G) mass as well as the absolute (H) and relative (I) epididymal fat mass were determined at the end of the experimental period. Absolute (D) and relative (E) epididymal fat mass as well as absolute (F) and relative (G) interscapular brown adipose tissue mass were determined at the end of the experimental period. The interscapular temperature (H–J) and the determination of interscapular brown adipose transcript expression of thermogenic genes were determined at the end of the experimental period. In all experiments, n=4–5; *p<0.05 vs. shNonTarget.

### Inhibition of Tril in POMC neurons of obese mice did not change the number of POMC neurons in the arcuate nucleus

In experimental obesity, there is increased apoptosis of neurons in the MBH, and this predominantly affects the number of POMC neurons (4–6). In obese mice submitted to the protocol of inhibition of Tril in POMC neurons (Fig. 7A), there were neither changes in the number of POMC neurons (Fig. 8A and 8B) nor in the cellular expression of cleaved caspase-3 (Fig. 8C and 8D).

**Figure 8.**
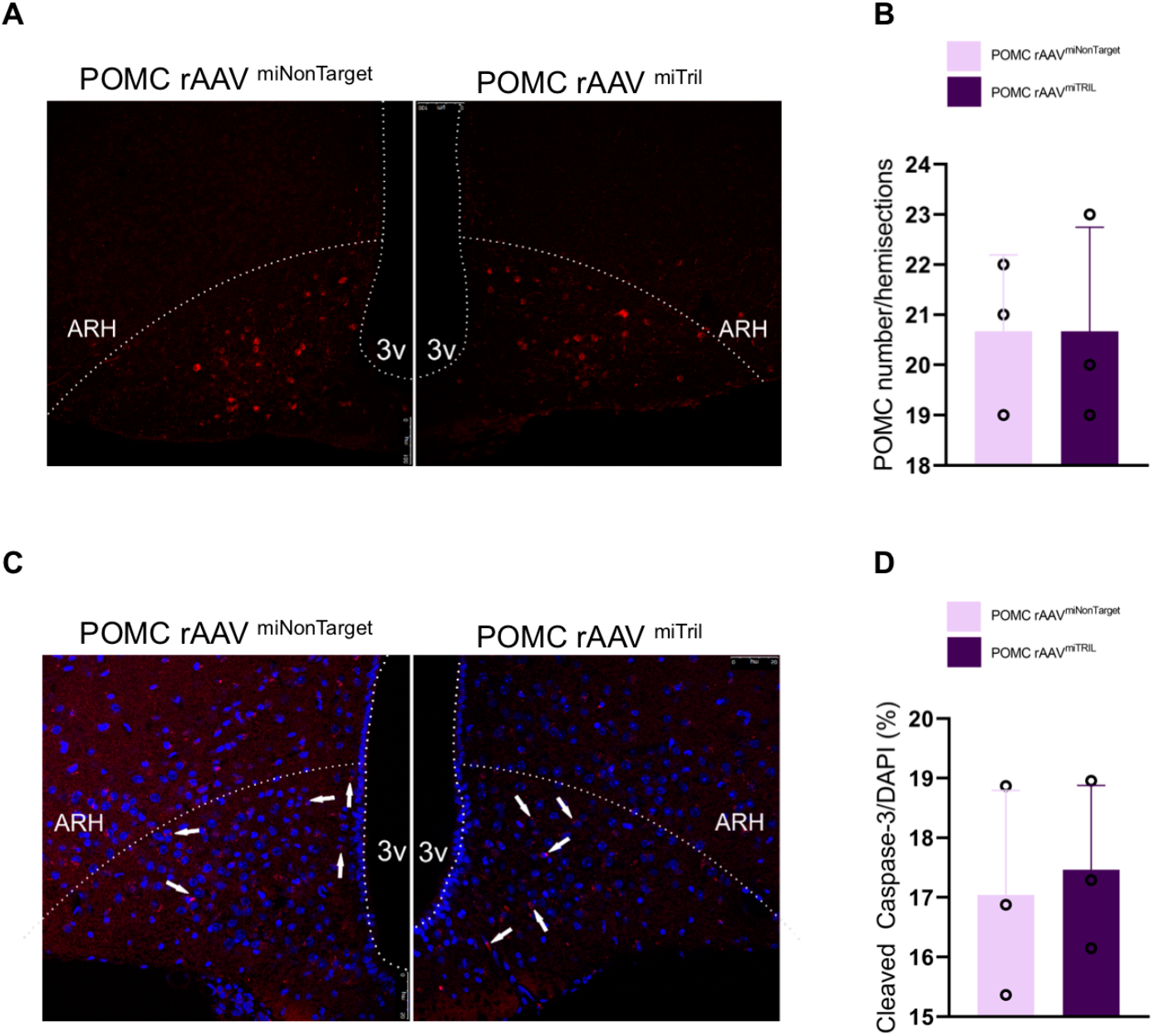
Inhibition of Tril in POMC neurons in mice submitted to long-term feeding on a high-fat diet fails to modify the number of POMC neurons and cells expressing cleaved caspase-3. POMC-Cre mice were fed a high-fat diet for 14 weeks and then submitted to an intracerebroventricular injection with Cre-dependent AAV-FLEX-EGFP-mir30 carrying either a non-target sequence (rAAVshNonTarget) or a Tril targeting sequence (rAAVshTril); after 2 weeks of recovery, mice were fed a high-fat diet for another 8 weeks (according to the protocol depicted in Fig. 7A). Immunofluorescence staining was employed to determine the number of cells expressing POMC (A and B) and cleaved caspase-3 (C and D). In all experiments, n=3. In C, arrows depict cells expressing cleaved caspase-3.

### Inhibition of Tril in the hypothalamus and POMC neurons has minor metabolic outcomes in mice fed chow

Most experiments described in the previous sections for mice fed a HFD were also performed in mice fed chow. Except for the changes in energy expenditure presented in Fig. 6A–6D, the inhibition of Tril in the hypothalamus, as well as specifically in POMC neurons, promoted no modification of metabolic parameters; therefore, we decided not to show the data.

## Discussion

The stability of body mass depends on the functional and structural fitness of MBH POMC neurons, as demonstrated by a number of different experimental approaches (11–14). In humans, mutations of MC4R result in the most common form of monogenic obesity (15, 16); in addition, in mutations of POMC, treatment with the MC4R agonist setmelanotide promotes sustainable reduction of hunger and weight (17), providing a clinical proof-of-concept for the central role of the hypothalamic melanocortin system in the regulation of body mass.

In DIO, POMC neurons are affected at the functional and structural levels by distinct but rather integrated mechanisms (1–3, 5, 18–20). The core mechanism behind DIO-associated POMC abnormalities is inflammation (21–25), and its drivers include the activation of RNA stress granules (26, 27), endoplasmic reticulum stress (2, 3, 19), PKCt (20) and TLR4 (2, 28). The long-term persistence of hypothalamic inflammation, as it occurs when mice are fed a HFD for longer than 8 weeks, leads to a progressive and potentially irreversible deterioration of the melanocortin system (4, 6, 23, 29–31). Interestingly, despite the fact that POMC and AgRP neurons share anatomical space, and thus, are potentially exposed to similar harmful factors, POMC undergoes more dramatic changes in response to DIO (4–6), raising the possibility that POMC-specific factors could be involved in this increased sensitivity to the diet.

Using a public dataset in a search for POMC-specific transcripts (8), we identified Tril, a TLR4-associated membrane protein previously shown to mediate the inflammatory response to LPS in the brain (9). Initially, we employed POMC and AgRP reporter mice to anatomically define the expression of Tril in the hypothalamus. We confirmed that Tril is mostly expressed in POMC neurons (8). In addition, as it could be expected for a TLR4-associated protein, Tril was detected in microglia (32). Using two distinct mouse strains, we showed that hypothalamic Tril is regulated by the consumption of a HFD, as other inflammatory proteins involved in the development of hypothalamic abnormalities in DIO (2, 3, 5, 33, 34).

In order to explore the potential involvement of hypothalamic Tril in the development of HFD-associated obese and metabolic phenotypes, we used two distinct approaches. First, we employed a lentivirus encoding a shRNA to selectively knockdown Tril in a non-cell-specific fashion in the arcuate nucleus (35). Second, we employed Cre-dependent adeno-associated viral vector to specifically inhibit Tril in POMC-expressing neurons (36). When either inhibition was performed in mice fed chow, there were virtually no changes in the phenotype, except for the increase in whole-body energy expenditure shown in Fig. 6A–6D that was obtained by specifically inhibiting Tril in POMC neurons. Conversely, when either inhibition was performed in mice fed a HFD, there were considerable beneficial changes in the obese and metabolic phenotypes. The different outcomes obtained in mice fed the distinct diets support the potential involvement of Tril in the diet-induced inflammatory response.

The action of Tril is similar to CD14 (serving as an accessory molecule that facilitates the binding of LPS to TLR4), mediating at least part of the inflammatory response triggered in this context (10). Studies have previously shown that Tril is expressed in glial cells (9, 10). However, as first shown by Henry and coworkers using RNA sequencing of hypothalamic neurons (8) and now, confirmed in this study, employing reporter mice, Tril is also expressed in POMC neurons. One important aspect of TLR4-associated protein expression in the hypothalamus is that, whether CD14 is expressed in glial cells, it is not expressed in POMC neurons (37); thus, in this particular cell population, Tril may be essential for the activation of TLR4.

The arcuate nucleus non-restricted and POMC-restricted Tril inhibitions generated similar but not completely overlapping phenotypes. Identity was found for reduction of body adiposity, increased whole body energy expenditure and reduction of BAT mass. In addition, there was a significant reduction of body mass gain in arcuate nucleus non-restricted Tril inhibition and trend for reduction in POMC-restricted Tril inhibition. As Tril is also expressed in glial cells (9, 10, 37), some of the effects obtained in the non-cell-specific inhibition might be related to the diet-induced activation of this protein in cells other than POMC cells.

Nevertheless, the importance of Tril in POMC neurons is evidenced by the robustness of the phenotype obtained when targeting was cell-specific. In other studies, inhibition of distinct components of the diet-induced responsive inflammatory machinery specifically in POMC neurons, such as Myd88, IKK/NFkB, HIF and TGF-βR (18, 26, 38, 39), resulted in beneficial changes in the metabolic phenotype placing POMC as a direct target for the harmful effects of a HFD. The results of our study place Tril as another important component of the complex mechanisms that affect POMC neurons in obesity.

In the final part of the study, we asked if the POMC-specific inhibition of Tril was capable of reverting the effects of long-term DIO. For that, mice were fed a HFD for 14 weeks before the inhibition of Tril in POMC neurons was undertaken. As previously shown, the metabolic outcomes of feeding mice a HFD for shorter than 8 weeks can be reverted by returning mice to chow; however, longer periods on a HFD lead to irreversible metabolic outcomes (31). In concert with human obesity, studies show that the longer patients remain obese, the more severe the clinical outcomes and the more resilient the obese phenotype is, irrespective of the therapeutic approach employed (40, 41). Here, the inhibition of Tril in POMC neurons in long-term obese mice resulted in no change in body mass, adiposity and food intake. There was only a trend to increase Ucp1 and a significant increase in Tmem26 in BAT, suggesting that, despite the lack of effect on the obese phenotype, the inhibition of Tril in long-term obese mice retains its thermogenic-inducing effect (42). In addition, the inhibition of Tril in POMC neurons in long-term obese mice was sufficient neither to change the number of arcuate nucleus POMC neurons nor the expression of cleaved-caspase 3.

In summary, Tril is a POMC-enriched protein that partially mediates the harmful effects of a HFD towards hypothalamic inflammation and systemic abnormal regulation of energy and glucose homeostasis. The capacity of regulating energy expenditure is particularly important as it occurred in all models and approaches tested in this study. The difference in POMC as compared to AgRP expression of Tril places this protein in a selected group of HFD-responsive proteins that are specific for POMC-expressing neurons.

## Methods

### Animal models

Six-week old, male, Swiss and C57BL/6J mice were obtained from the University of Campinas experimental animal breeding facility. AgRP-IRES-Cre mice (Agrptm1(cre)Lowl/J, Jackson Laboratories) were crossed with the Cre-inducible tdTomato reporter mouse (B6;129S6-Gt(ROSA)26Sortm9(CAG-tdTomato)Hze/J, Jackson Laboratories), and POMC-Cre were used in Cre-dependent rAAV experiments or crossed with Cre-inducible GFP-reporter mice (The Jackson Laboratory) to determine the colocalization of Tril with fluorescently labeled AgRP and POMC neurons, respectively. Mice were housed individually at 22°C (±1°C) using a 12 h light/12 h dark cycle. All mice had ad libitum access to chow (3.7 kcal g^−1^) or a HFD (5.1 kcal g^−1^) and water. For some experiments, mice were submitted to periods of fasting, as described elsewhere in the text. All experiments were approved by the Ethics Committee at the University of Campinas (#4069-1 and #4985-1).

### Respirometry

To determine O_2_ consumption, CO_2_ production and energy expenditure, mice were acclimatized for 48 h in an open circuit calorimeter system, the LE 405 Gas Analyzer (Panlab – Harvard Apparatus, Holliston, MA, USA). Thereafter, data were recorded for 24 h. Results are presented as the average of light and dark cycles.

### Glucose and insulin tolerance tests

Following 6 h of fasting, mice received intraperitoneal (ip) injections with solutions containing glucose (2.0 g/kg body weight) or insulin (1.5 IU/kg body weight) and then blood samples were collected for ip-glucose tolerance test (ipGTT) or ip-insulin tolerance test (ipITT), respectively. Glucose concentrations were measured in tail blood using a portable glucose meter (Optium Xceed, Abbott) at 0, 15, 30, 60 and 120 minutes after glucose administration or 0, 5, 10, 15, 20, 25 and 30 minutes after insulin administration.

### Leptin responsiveness

Mice were submitted to 24 h of fasting and then received ip injections with either saline (0.9% NaCl) or mouse recombinant leptin (5 mg/kg body weight, Calbiochem, Billerica, MA, USA) immediately before the dark cycle; food intake was recorded 1, 2, 4, 8 and 12 h following the injection.

### Assessment of body composition

To determine total body fat and lean mass, time-domain nuclear magnetic resonance (TD-NMR) was applied using the LF50 body composition mice analyzer (Bruker, Germany). Measurements were performed on the last day of the experiment.

### Thermal images

The estimated iBAT temperatures were determined using an infrared (IR) camera (FLIR T450sc, FLIR systems, Inc. Wilsonville, USA) and analyzed with FLIR-Tools software.

### Tissue collection and histology

C57BL/6J, AgRP tdTomato and POMC-GFP mice were deeply anesthetized with ketamine (100 mg/kg) and xylazine (10 mg/kg) and submitted to transcardiac perfusion with 0.9% saline followed by 4% paraformaldehyde (PFA). Brains were removed, postfixed 24 h in 4% PFA solution and then transferred to a solution containing 20% sucrose in 0.1 M PBS (pH 7.4) for 12 h. Perfused brains were frozen at −30°C and sectioned on a cryostat at a thickness of 30 μm. For immunohistochemistry, free-floating sections were washed three times for 10 minutes with 0.1 M PBS. Next, sections were blocked in 5% donkey serum and 0.2% Triton X-100 in 0.1 M PBS for 1 h at room temperature, followed by incubation in goat anti-TRIL primary antibody (sc-24489, Santa Cruz Biotechnology, Santa Cruz, CA, USA), rabbit anti-IBA1 (Wako Chemicals, #019-19741), rabbit anti-cleaved caspase-3 (Cell Signaling, #9661S), rabbit anti-POMC (Phoenix Pharmaceuticals, #H-029-30), sheep anti-αMSH (Chemicon, #AB5087) and rabbit anti-AgRP (Phoenix Pharmaceuticals, #H-003-53) for 24 h. The sections were then washed three times in 0.1 M PBS and incubated for 2 h at room temperature in donkey anti-goat AlexaFluor^546^ (1:500, Invitrogen, #A-11056), donkey anti-goat FITC (1:500, Santa Cruz, sc-2025), donkey anti-rabbit FITC (1:500, Abcam, ab6798), donkey anti-rabbit IgG AlexaFluor^594^ (1:500, Jackson Immuno Research, #711-585-152) and donkey anti-sheep AlexaFluor^488^ (1:500, Jackson Immuno Research, #713-545-003) conjugated secondary antibodies. Thereafter, the sections were mounted onto slides, and the nuclei were labeled with TOPRO (Life Technologies, T3605). The sections were analyzed with a LEICA TCS SP5 II confocal laser-scanning microscope (Leica Microsystems, Wetzlar, Germany).

### pSTAT3 staining

Mice were submitted to 12 h of fasting and then received ip injections with mouse leptin (5 mg/kg body weight, Calbiochem, Billerica, MA, USA). After 1 h, mice were submitted to transcardiac perfusion with 0.9% saline followed by 4% PFA. After sectioning on a microtome, the brain sections were rinsed in 0.02 M KPBS (pH 7.4), followed by pretreatment in a water solution containing 1% hydrogen peroxide and 1% sodium hydroxide for 20 minutes. After extensive washings in 0.02 M KPBS, the sections were incubated in 0.3% glycine for 10 minutes and then 0.03% lauryl sulfate for 10 minutes.

Thereafter, the sections were blocked in 3% normal donkey serum for 1 h, followed by incubation in rabbit anti-pSTAT3^Tyr705^ (1:1000, Cell Signaling, #91315) for 48 h. For the immunofluorescence reactions, sections were rinsed in KPBS and incubated for 120 minutes in Fab fragment donkey anti-rabbit AlexaFluor^488^ (1:500, Jackson Immuno Research, #711-547-003). Thereafter, sections were washed three times in 5% formalin for 10 minutes and then washed three times in 0.02 M KPBS for 5 minutes. Sections were then incubated with POMC antibody (1:1000. Phoenix Pharmaceuticals, #H-029-30) overnight at room temperature. After washing in 0.1 M PBS, the sections were incubated for 30 minutes with the secondary antibody AlexaFluor^594^ (1:500, Jackson Immuno Research, #711-585-152) diluted in 0.02 M KPBS. After three washes in 0.02 M KPBS for 10 minutes, the sections were mounted onto gelatin-coated slides and coverslipped with Fluoromount G (Electron Microscopic Sciences, Hatfield, PA, USA). The percentage of pSTAT3-positive POMC neurons was determined in blind counting by three distinct researchers using hemisections of the middle ARH from four animals for statistical comparison. The sections were processed simultaneously under identical conditions and analyzed with the same microscope set-up.

### Lentiviral clones

Three different shRNA clones targeting Tril (Sigma-Aldrich, St Louis, MO, USA) and scramble lentiviral particles were used for the overall knockdown experiments (35). For arcuate nucleus (Arc) bilateral lentiviral delivery, 8–12 week old male C57BL/6J mice were anesthetized with ketamine (100 mg/kg body weight) and xylazine (10 mg/kg body weight) intraperitoneally, and the stereotaxic surgery was carried out using a stereotaxic frame (Stoelting Apparatus, Wood Dale, IL, USA) set at AP −1.7 mm, ML ±0.3 mm and DV −5.6 mm coordinates from Bregma.

### Cre-dependent recombinant adeno-associated viral (rAAV) vectors

With regard to the POMC-specific knockdown of Tril, referred to here as POMC rAAV^miTRIL^, a TRIL-based miRNA construct was constructed by modifying the Cre-dependent AAV-FLEX-EGFP-mir30 (Scn9a) (Addgene plasmid # 79672) (43, 44). Briefly, the plasmid was modified by the insertion of an shRNA sequence against TRIL between *Eco*RI and *Xho*l. The sequence used was the following, where complementary sequences are underlined: 5’CGAGGCAGTAGGCA<u>CCAGTATCTTACTGTGTTATT</u>TACATCTGTGGCTTCACTA<u>AATAACACAGTAAGATACTGG</u>CGCTCACTGTCAACAGCAATATACCTT-3’.

The rAAV particles were produced in the LNBio (Brazilian Biosciences National Laboratory) facility and titrated using an Addgene protocol by qPCR (45). Briefly, purified AAV particles were treated with DNAse by incubation for 30 min at 37°C. A standard curve was generated using the AAV-FLEX-EGFP-mir30 (TRIL) plasmid serial diluted. SYBR Green quantitative PCR (qPCR) was performed using the primers: fwd ITR primer 5’-GGAACCCCTAGTGATGGAGTT-3’ and rev ITR primer 5’-CGGCCTCAGTGAGCGA-3’. The Cre-dependent bilateral injections of the rAAV vectors in the ARH were performed in POMC-Cre mice at the following stereotaxic coordinates: AP −1.7 mm, ML ±0.3 mm and DV −5.6 mm from Bregma. All AAVs were allowed 2 weeks for expression before experiments were initiated.

### Gene expression analysis

Total RNA was extracted from the hypothalamus, inguinal and epididymal white adipose tissue, brown adipose tissue and liver using TRIzol reagent (Invitrogen). cDNA synthesis was performed using 2 μg of total RNA. The PCR containing 25 ng of reverse-transcribed RNA was performed using the ABI Prism 7500 sequence detection system (Applied Biosystems). For RT-PCR calculation, the delta CT was used, and the relative gene expression was normalized to that of GAPDH in all samples. The primers used were Tril (Mm01330899_s1), Ucp1 (Mm01244861_m1), PGC1α (Mm01188700_m1), Zic1 (Mm00656094_m1), LHX8 (Mm00802919_m1), Tfam (Mm00447485_m1), PPARγ (Mm01184322_m1), Scd1 (Mm00772290_m1), Srebf1 (Mm01138344_m1), Cd36 (Mm01135198_m1, Tnfα (Mm00443258_m1), F4/80 (Mm00802529_m1), Il1-β (Mm00434228_m1), Nlrp3 (Mm00840904_m1), Hsp90 (Mm00441926_m1), Hspa5 (Mm00517691_m1), Caspase-3 (Mm01195085_m1), Pomc (Mm00435874_m1), AgRP (Mm00475829_g1), Mch (Mm01242886_g1), Tmem26 (Mm01173641_m1), Th (Mm00447557_m1), Ccl2 (Mm99999056_m1) and Tbx1 (Mm00448949_m1).

### Western blotting

Hypothalamic specimens were homogenized in solubilization buffer (1% Triton X-100, 100 mM Tris (pH 7.4), 100 mM sodium 22 pyrophosphate, 100 mM 4 sodium fluoride, 10 mM EDTA, 10 mM sodium vanadate, 2 mM 23 PMSF and 0.1 mg/mL 5 aprotinin). A total of 100 μg of protein per sample were separated by sodium dodecyl sulfate-polyacrylamide gel electrophoresis (SDS-PAGE), transferred to nitrocellulose membranes and blocked in 3% BSA solution in TBST for 2 h. After a washing step, the membranes were blotted with goat-anti TRIL antibody (sc-24489, Santa Cruz Biotechnology, Santa Cruz, CA, USA), and α-tubulin (Sigma-Aldrich, T5168) was used as loading control.

Specific bands were labeled by chemiluminescence and were quantified by optical densitometry after exposure to Image Quant LAS4000 (GE Healthcare, Life Sciences).

### Statistical analysis

Results are presented as mean ± standard error of the mean (SEM). Statistical comparisons between different times of refeeding and controls were performed using analysis of variance (ANOVA), followed by Tukey’s post hoc test. Student’s t tests were applied for comparisons between scramble and KD^TRIL^ in experiments with lentiviral clones or POMC Cre^miNonTarget^ and POMC Cre^miTRIL^ in experiments with Cre-dependent rAAV.

## Author Contributions

AM-A and LAV conceptualized and designed the experiments; AM-A, performed the majority of the experiments; PAN, JCLJr and JMG contributed to mouse experiments and analytical methods; FMS designed, cloned and titrated the Cre-dependent rAAV; JDJr contributed to histological analyzes and provided POMC-GFP, AgRP tdTomato and POMC Cre mice; AM-A and LAV discussed and organized results; AM-A and LAV wrote the paper; LAV was also responsible for funding acquisition. All authors contributed to the editing and discussion of the manuscript.

## Acknowledgments

Alexande Moura-Assis received financial support from the São Paulo Research Foundation (FAPESP #2016/01245-5). Pedro A. Nogueira received financial support from Coordenadoria de Aperfeicoamento de Pessoal de Nivel Superior through the graduate Course in Cellular and Structural Biology, Insititute of Biology, University of Campinas. The authors thank Erika Roman, Joseane Morari, Marcio Cruz and Gerson Ferraz for laboratory management.

## Funding

LAV, São Paulo Research Foundation (FAPESP – 2013/07607-8). FMS, São Paulo Research Foundation (FAPESP – 2018/14818-9). JDJr, São Paulo Research Foundation (FAPESP – 2017/02983-2). Conselho Nacional de Desenvolvimento Cientifico e Tecnológico, INCT-Neuroimmunomodulation. Coordenadoria de Aperfeiçoamento de Pessoal de Nível Superior

## Notes

**Conflict of interests:** The authors declare that no conflicts of interests exists.

### Competing Interest Statement

The authors have declared no competing interest.

